# Scalable microvascular networks-on-chip enable 8-week unidirectional perfusion for long-term vascular toxicity screening

**DOI:** 10.64898/2026.06.10.731320

**Authors:** A Rodrigues, SPM Ruiter, I Schilt, C Clavijo, T Olivier, P Vulto, TP Burton, LJ van den Broek

**Affiliations:** MIMETAS BV, Oegstgeest, The Netherlands; Leiden Academic Centre for Drug Research, Leiden University, Leiden, The Netherlands

**Author notes:** These authors jointly supervised this work (TP Burton, LJ van den Broek).

**Keywords:** Vascular networks, Organ-on-a-chip, Unidirectional flow, long-term stable, acute and chronic toxicity

## Abstract

Cardiovascular toxicity is a leading cause of late-stage drug attrition, yet current *in vitro* vascular models lack the longevity and scalability needed to capture clinically relevant responses in mature vasculature. We developed a vascular microphysiological system comprising 32 parallel, self-assembled microvascular networks from primary human endothelial cells and stromal fibroblasts. Networks were unidirectionally perfused by gravity-driven, pump-free flow. They remained functionally perfusable for at least 57 days, the longest duration reported for self-assembled microvascular networks, and underwent progressive maturation characterised by perivascular fibroblast organisation, basement membrane deposition, and matrix remodelling. Variance decomposition confirmed high reproducibility, with intra-plate variability of 3–7% and no operator-dependent effects on network morphometry. Exploiting this extended culture window, we reveal maturation-dependent shifts in endothelial inflammatory responsiveness and cytotoxic susceptibility. Distinct acute and chronic toxicity profiles of clinically used tyrosine kinase inhibitors are resolved, and a concentration- and time-dependent spectrum of sorafenib-induced microvascular responses is produced. These results show that vascular longevity and progressive maturation are key requirements for assessing vascular toxicity, and provide a scalable platform for evaluating acute and chronic drug responses in mature human microvascular tissues.

## INTRODUCTION

The human microvasculature plays a central role in tissue homeostasis by regulating oxygen and nutrient exchange, immune surveillance, and inflammatory signalling across organ systems^1^. Disruption of vascular integrity and endothelial barrier function contributes to the pathogenesis of numerous chronic diseases, including cardiovascular disease, cancer, fibrosis, retinal disorders, autoimmune disorders and diabetes-associated complications^2,3^. Moreover, acute and chronic vascular toxicities are among the top three causes of late-stage drug attrition and post-marketing safety withdrawals^4,5^. Experimental systems capable of reproducing human vascular physiology are therefore essential for understanding disease mechanisms and evaluating therapeutic responses.

Historically, animal models have been used to study vasculature in the context of disease and drug response. However, such models are inherently non-human. Cell culture techniques are undergoing rapid innovation, from largely cell line-based monocultures to full-fledged organoid models that recapitulate embryonic or regenerative development of organs^6^. In a surge towards ever more comprehensive cultures, increased efforts are focusing on integrating vasculature into iPSC-derived organoids, or through co-culture approaches in adult stem cell (ASC) organoids or spheroids^7^. Although such models have yielded reasonable success, long-term stable vasculature has not been achieved, mainly due to a lack of perfusion^8^.

Microphysiological systems (MPS) add perfusion flow as a critical component to endothelial cell culture, with meaningful progress reported ^9,10^. However, incorporating relevant architecture and longevity in vascular MPS is yet a scarcely explored aspect^11^. Many published systems either lack stromal-vascular co-culture, limiting paracrine crosstalk, or rely on synthetic membranes that physically separate cellular compartments, constraining cell-matrix contact and juxtacrine signaling^12,13^. Conversely, platforms supporting self-assembled microvascular networks (SAMNs) address some of these biological limitations by enabling multicellular, three-dimensional architectures; however, they often involve technically demanding setups that compromise scalability and reproducibility. Furthermore, many of these systems struggle to maintain long-term functional and perfusable vascular networks beyond 14 days, thereby constraining compound-testing studies to early-stage models or *de novo* angiogenesis rather than mature networks ^14–16^. This represents a critical bottleneck as many clinically relevant compounds, including tyrosine kinase inhibitors (TKIs) such as sorafenib, also act on established microvasculature and induce acute as well as chronic vascular responses. Importantly, effects on barrier function, perfusion, and vascular toxicity may occur independently of angiogenic activity and therefore require experimental systems capable of maintaining mature, functional microvascular networks over extended periods^17–19^

Physiological perfusion represents an additional requirement for modelling vascular function *in vitro* ^20^. Unidirectional flow regulates endothelial alignment, strengthens intercellular junctions, and promotes vascular homeostasis, whereas disturbed or oscillatory flow drives endothelial dysfunction and transcriptional reprogramming *in vivo*^21,22^. Yet, many vascular MPS rely on pump-based perfusion systems that introduce tubing complexity, pulsatility, and risks of bubble trapping, thereby complicating workflows, increasing operational costs, and limiting throughput. Pump-free systems that incorporate unidirectional flow represent a promising yet underexplored path forward, combining hemodynamic relevance with operational simplicity.

Recent studies have addressed individual aspects of these challenges. Ehlers et al. made important advancements by implementing pump-free, gravity-driven unidirectional flow in a microtiter-plate format, confirming endothelial alignment and maintenance of a non-contractile smooth muscle phenotype^23^. However, the system is limited to a single endothelial tube and is therefore not suitable for studying microvasculature remodelling. Similarly, a brain microvasculature platform by Admiraal et al. utilized partially unidirectional flow to support perfusability, yet did not demonstrate long-term viability^24^. To address longevity in SAMNs, Floryan et al. developed a PDMS-based system demonstrating that unidirectional flow maintained long-term network perfusability; however, this system relied on immortalized endothelial cells and required an external pump for inducing unidirectional flow.^25^. Therefore, no platform has yet combined self-assembled microvascular complexity with long-term perfusability in a scalable, robust and reproducible manner.

In the current study, we aim to address this gap with a novel, scalable platform engineered to support complex vascular network formation - the OrganoPlate® Graft 32 UniFlow (UF). This system addresses key technical and translational challenges by combining a membrane-free culture configuration in an open-well format with the simplicity of gravity-driven unidirectional perfusion, while preserving scalability.

We demonstrate that this platform consistently generates 32 perfusable vascular networks in parallel with high reproducibility and minimal variability. Supported by stromal cells and ECM, three-dimensional (3D) microvascular networks self-assemble and maintain functional perfusion and barrier integrity for at least 57 days. We examine inflammatory immune cell responses, acute compound-induced vascular and stromal toxicity, and vascular responses to chronic drug exposure. Importantly, the phenotypic endpoints employed – including barrier integrity, perfusability and network morphometry – provide phenotypic, non-invasive assessment that allow to follow compound effects over prolonged periods of time. Collectively, these findings present a robust and scalable vascular MPS that advances the current standard for *in vitro* vascular modelling. As such, it is well-positioned to support vascular safety screening, cardiotoxicity assessment, and the study of inflammatory vascular diseases throughout drug discovery and development workflows.

## RESULTS

### Self-assembled microvascular networks can be generated under gravity-induced unidirectional perfusion

To enable the generation of physiologically relevant, reproducible perfusable microvascular networks, we developed a novel microfluidic platform: the OrganoPlate® Graft 32 UF (Fig. 1). The OrganoPlate platform is built on a microtiter plate footprint, comprising 32 independent chips underneath the titerplate. An evaporation rim around the well-grid is filled with sterile PBS to prevent edge effects due to humidity gradients. The chips comprise #1 coverslip-thickness glass for plate flatness, optimal optical clarity, enabling high-content high magnification imaging (Fig. 1a). Each individual chip comprised two main fluidic circuits — a perfusion channel, a bypass channel and a gel chamber (Fig. 1b). The chip layout is formed from a 6 × 2 well arrangement, in which the top six wells are merged to form two reservoirs (left and right merged wells) that contain the inlet–outlet interfaces for both perfusion and bypass channels. The gel chamber features an open-top configuration enabling direct chamber filling with extracellular matrix (ECM). Surface-tension-based ECM patterning using phaseguide technology enables a layered culture, while eliminating the need for artificial membranes^26^.

**Figure 1.**
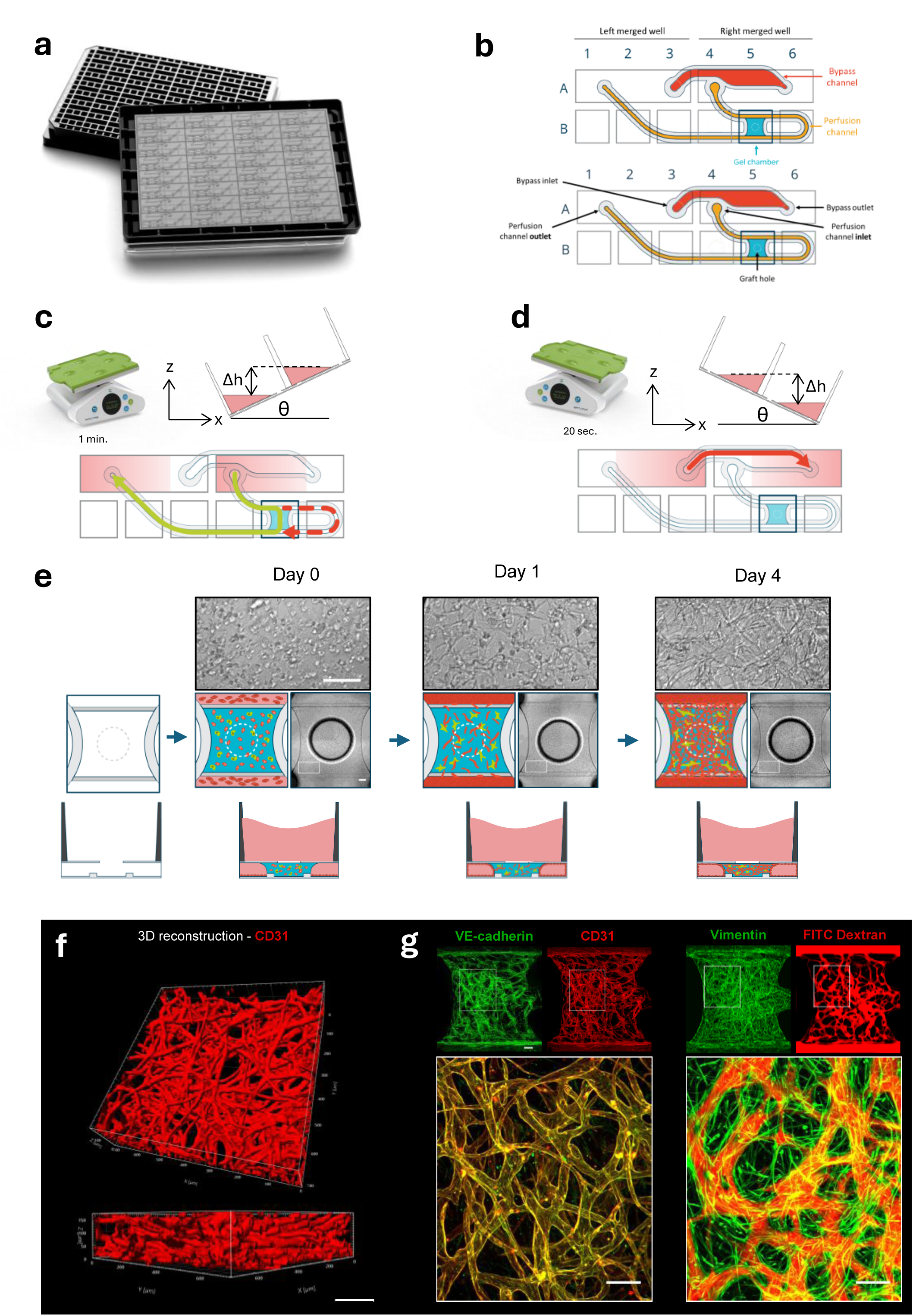
| The OrganoPlate Graft 32 UF supports gravity-driven unidirectional flow through a three-dimensional self-assembled microvascular network. **a,** The OrganoPlate Graft 32 UF is a modified 384-well plate with a glass bottom integrating 32 individual microfluidic chips, each supporting gravity-driven, unidirectional flow through a self-assembled microvascular network. **b,** Schematic of a single microfluidic chip in a 6x2 well configuration, comprising a central gel chamber (blue) with a 1 mm diameter access port for loading an ECM-embedded cell suspension, a perfusion channel (orange), and a bypass channel (red). **c,d,** OrganoPlates are positioned on a rocking platform to generate gravity-driven, unidirectional flow. During the positive incline (c), medium enters through the inlet into the perfusion channel and flows through the (once formed) vascular network in the gel chamber. During the negative incline (d), fluid is redirected through the bypass channel, resetting the system. **e,** Representative schematics of the top and side view of the gel chamber from the OrganoPlate Graft 32 UF. HUVECs (red) and NHLFs (green) were embedded in a fibrin-based ECM (blue) and loaded into the gel chamber. HUVECs were subsequently seeded into the perfusion channel to allow tubule formation. Close-up phase-contrast images on days 0, 1, and 4 illustrate progressive self-assembly into an interconnected vasculature. Scale bar, 200 µm. **f,** 3D reconstruction of immunofluorescent CD31-positive structures at day 15 demonstrates formation of an intricate network. Multiple z-planes (1µm step-size) acquired within a single site at the center of the gel chamber with a 20X objective. Scale bar, 150 µm. **g,** At day 15, immunofluorescent staining for CD31 and VE-cadherin illustrates endothelial vessel architecture. Vimentin staining reveals a stromal fibroblast network closely associated with the vasculature. Perfusion with fluorescent dextran confirmed vessel functionality and interconnectivity throughout the network. Entire network (complete gel chamber) results from a 4-site acquisition per chip with a 10X objective. Close-up images were generated using Fiji from the overlay. Scale bar, 200 µm. Images are representative of n≥20 chips from N=3 independent experiments.

To validate the platform vascular network formation, we established a co-culture of human umbilical vein endothelial cells (HUVECs) and normal human lung fibroblasts (NHLFs) mixed with a fibrin ECM and loaded into the gel chamber (Fig. 1c). The cell-laden fibrinogen-thrombin suspension was introduced via the top opening into the gel chamber. Two microfabricated phaseguides that act as capillary pressure barriers pinned the suspension, thus preventing overflow into adjacent lanes. Following ECM polymerisation, HUVECs were seeded into the fibronectin-coated perfusion channel, allowing the formation of a continuous endothelial tubule along the channel interface. By day 1, endothelial cells underwent a morphological transition from amoeboid to elongated, vasculogenesis commenced within the ECM and a confluent endothelial tubule was established in the perfusion channel. By day 4, an intricate and dense culture was evident in phase contrast images of the gel chamber, with endothelial network branches and fibroblasts homogeneously covering the entire gel and connecting with the HUVEC tubules in the adjacent perfusion channel (Fig. 1e).

Continuous medium perfusion through the vascular network was achieved by gravity-driven flow induced via plate rocking (Fig. 1c, d). Inclination of the OrganoPlate® Graft 32 UF under a first, negative, angle creates a hydrostatic pressure gradient along the length of the perfusion channel, inducing flow across the microvascular bed during. In this active phase, medium flows through the perfusion channel, this create a pressure gradient across the gel chamber which drives flow through the ECM-embedded microvascular bed once connections have been formed (Fig. 1c). The system is reset through the bypass channel during inclination under a second, positive, angle (Fig. 1d). Through continuous rocking, effective recirculation of growth medium is achieved, sustaining unidirectional perfusion across the network, and ensuring nutrient and oxygen exchange throughout the culture period. Uninterrupted unidirectional perfusion through the parent vessel and across the vascular network is induced for 60 seconds per tilt-cycle, with limited transient backflow (0-8 seconds) during rocker transitions and a 20 second reset phase (depending on the self-organising microvascular connectivity heterogeneities per chip). The flow profile and shear stress experienced by both the parent vessel and microvasculature of each chip is a function of the number of connections made and width of the vessels within the network.

Structural and functional maturity of the engineered microvasculature was confirmed by immunofluorescent staining and perfusion assays at day 15 (Fig. 1f, g). Confocal imaging revealed an intricate 3D vascular network that stained positive for the endothelial marker CD31, extending throughout at least 70% of the gel chamber height (z > 170 µm) (Fig. 1f). Additionally, VE-cadherin was localized to inter-cellular junctions along vessel walls and confirmed intact endothelial junctions and luminal continuity.

In parallel, vimentin staining revealed an extensive stromal network of fibroblasts within the gel chamber. Fluorescent dextran perfusion confirmed that the vessels were functional and interconnected, with tracer filling throughout the network. A close-up image shows vimentin-positive fibroblasts associated with perfusable vessels (Fig. 1g), suggesting stromal-endothelial interactions.

Together, these results show that the platform supports the formation of multicellular microvascular networks under unidirectional perfusion within a scalable, imaging-compatible format.

### Microvascular networks can be robustly and reproducibly generated across experiments and operators

After demonstrating the platform’s capacity to generate intricate microvascular networks, we evaluated its robustness by assessing intra-plate variability and spatial uniformity across a 32-chip plate (Fig. 2a). To assess potential "edge effects" common in cell culture including microfluidic platforms, we mapped the perfusable network area fraction across the plate using a spatial heatmap. The maximum deviation from the mean perfusable area fraction was restricted to 8.84% (Fig. 2a), with no apparent association between chip location and perfusable area as shown by the lack of statistical significance between edge and interior chips (Supplementary Fig.1). This high level of intra-plate consistency is further supported by low coefficients of variation (%CV) across independent plates (4.776% and 4.953% for plates 1 and 2, respectively; Supplementary Table 1), highlighting the platform’s capacity for consistently generating vascular beds.

**Figure 2.**
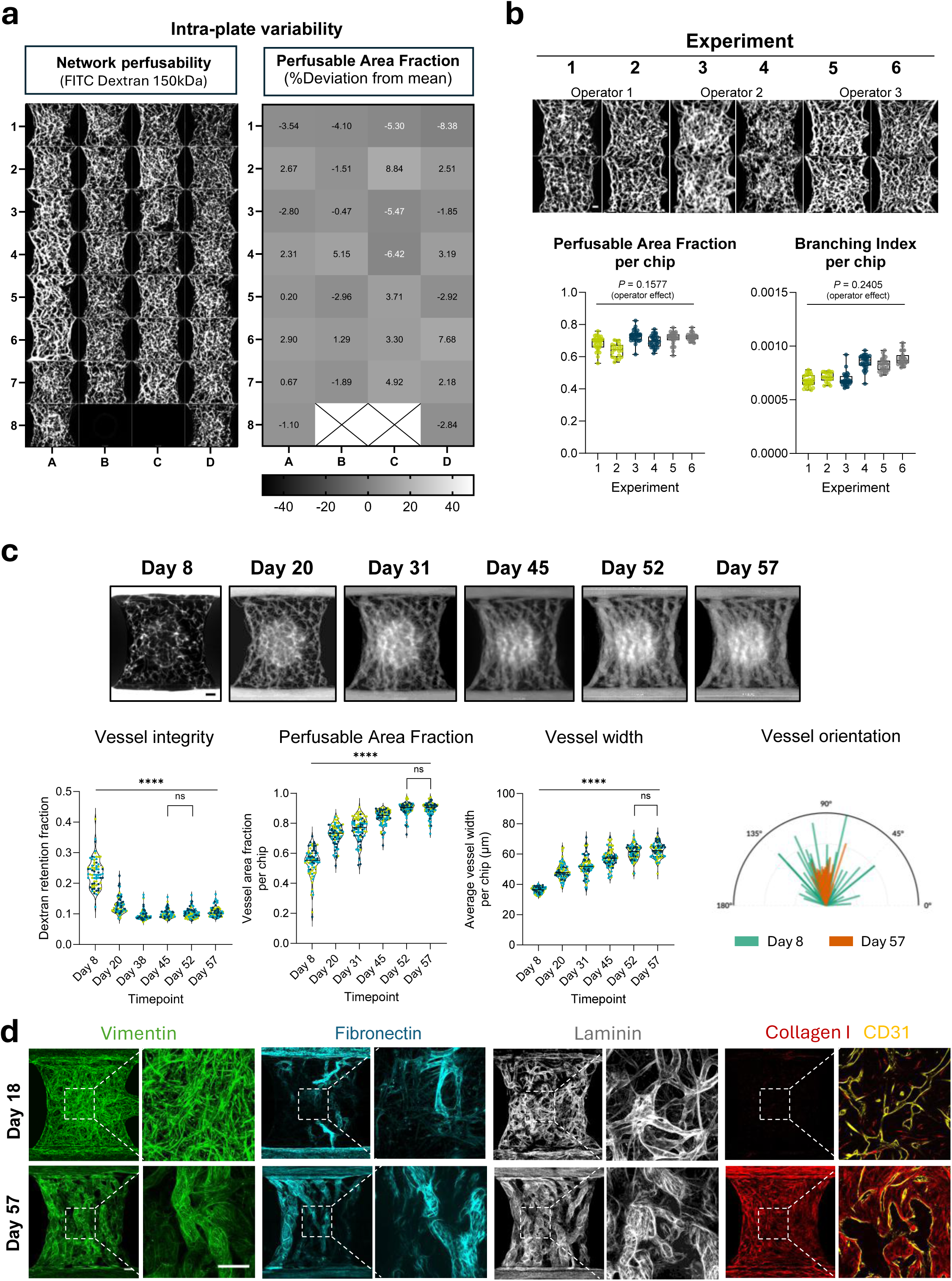
| Microvascular networks exhibit stable and robust maturation over 57 days of culture. a, Intra-plate variability in microvascular network organisation was assessed by perfusing cultures with 150 kDa fluorescein isothiocyanate (FITC)-dextran at day 20. Perfusable area fraction was quantified and expressed as percent deviation from the plate mean (n=30 chips per plate). Chips 8B and 8C represent cell-free controls. **b,** Representative fluorescence images of dye perfusion assay from microvascular networks prepared by three independent operators across six independent experiments. Scale bar, 200 µm. Perfusable area fraction and branching index were quantified across all six experimental batches. Each data point represents an individual chip (N=6 independent experiments, n=172 chips total). Colours indicate operator identity. Nested one-way ANOVA revealed no significant operator effect for either parameter; significant variability was detected between experiments nested within operators (P<0.0001), consistent with expected biological replicate variability. **c,** Representative fluorescence images of dye perfusion assay in microvascular networks at six timepoints over 57 days of culture (4X objective). Scale bar, 200 µm. Vessel integrity assessed via 150 kDa FITC-dextran retention fraction, perfusable area fraction, and average vessel width were quantified at each timepoint. Each data point represents a single chip; colours indicate independent experimental plates. For vessel integrity, N=3 independent experiments, n=75 chips per timepoint; for perfusable area fraction and vessel width, N=3, n=85 chips per timepoint. Overall effect assessed by repeated measures one-way ANOVA; only non-significant (ns) comparisons are indicated. Exact P values for all pairwise comparisons are provided in Supplementary Tables 5, 6 and 7. Brackets indicate non-significant comparisons between adjacent timepoints (Dunnett’s post hoc test); all other comparisons were statistically significant. Vessel orientation was quantified at two timepoints; each needle represents a single chip, where needle length corresponds to the directionality magnitude (sum of the histogram from centre − SD to centre + SD divided by the total histogram sum) and angle corresponds to the Gaussian centre (N=2 independent experiments, n=27–32 chips per plate). **d,** Immunofluorescent staining at day 18 and day 57 for vimentin, fibronectin, laminin (all maximum intensity projections), collagen I, and CD31 (single z-plane) illustrating remodelling of the basement membrane and interstitial matrix over time. Full network images represent a 4-site acquisition per chip using a 10X objective; scale bar, 200 µm. Higher magnification images represent single-site acquisition using a 20X objective; scale bar, 200 µm. Images in b–d are representative of the stated N and n. Significance thresholds: *P<0.05, **P<0.01, ***P<0.001, ****P<0.0001.

To assess inter-experimental reproducibility, nested one-way ANOVA analyses were performed for both area fraction and branching index across three independent operators and six experimental batches (n = 172 per parameter) (Fig. 2b). No significant operator-dependent effects were detected for either area fraction (P = 0.158) or branching index (P = 0.24), demonstrating robustness of the protocol to personnel-associated variability. Although statistically significant inter-experiment effects were observed for both parameters (P < 0.0001), the associated effect sizes remained small relative to the intrinsic chip-to-chip variability. For area fraction, the variance attributed to batch-to-batch differences (0.00048) was approximately 2.9-fold lower than the intra-experimental chip-level variance (0.00138), which accounted for more than 74% of the total random variation. Similarly, for branching index, inter-batch variance (5.22 × 10⁻⁹) remained within the same order of magnitude as the baseline chip-level variance (3.395 × 10⁻⁹). Consistent with these findings, intra-experimental precision remained high across all experiments, with coefficients of variation ranging from 3.0-6.5% for area fraction and 6.3-8.9% for branching index (Supplementary Table 2). Collectively, these results demonstrate a high degree of reproducibility and experimental robustness of the model across operators and independent experimental runs.

### Microvascular networks remain perfusable for 57 days and undergo maturation

To assess whether the networks could sustain perfusion over extended culture periods, we tracked the evolution of the microvasculature over six timepoints using longitudinal dextran perfusion across three independent plates. These analyses confirmed the functional longevity of the platform, with the networks remaining fully perfusable for at least 57 days (Fig. 2c).

Morphometric analysis (Fig. 2c) revealed progressive structural remodelling across the culture period, with vessel area fraction and average vessel width both increasing significantly over time (RM one-way ANOVA, P < 0.0001; n = 82 per timepoint), reaching a plateau by Day 52 with no further significant changes detected through Day 57. Quantification of network integrity via 150 kDa dextran retention fraction revealed a significant, time-dependent maturation of the endothelial barrier (P < 0.0001). Following a period of initial developmental variability at Day 8, macromolecular retention remained consistently low from Day 20 to Day 57, demonstrating sustained barrier function throughout the full period of active vascular remodelling.

To investigate the influence of unidirectional flow on network morphology, vessel orientation was analysed at Day 8 and Day 57 (Fig. 2c). At Day 8, vessels exhibited a broad, isotropic angular distribution with no preferred orientation. By Day 57, the network showed an apparent shift toward alignment at 90°, consistent with the axis of applied flow. These findings demonstrate that the platform supports long-term vascular maturation combining structural remodelling, sustained barrier function, and flow-driven network organization.

To characterise the biological mechanisms underlying this long-term stability, we compared the evolution of the perivascular niche and ECM between day 18 and day 57 (Fig. 2d). Immunostaining for vimentin revealed a transition from loose fibroblast distribution at day 18 to structured perivascular layers encasing the vessels at day 57 consistent with a pericyte-like stabilising role. This cellular reorganisation coincided with remodelling of the basement membrane and interstitial matrix. Laminin transitioned from a diffuse lining at day 18 to discrete perivascular bundles at day 57, indicative of progressive basement membrane maturation (Fig. 2d). Fibronectin persisted in selected vessels at Day 57, consistent with incomplete replacement of the provisional matrix, while collagen I appeared more extensively distributed within the interstitial spaces, potentially reflecting gradual remodelling of the initial fibrin-based scaffold.

### Vascular maturation state impacts dynamics of immune cell adhesion

To evaluate the functional capacity of the platform to recapitulate the human inflammatory cascade, we investigated the recruitment and extravasation of PBMCs within the established microvascular networks (Fig. 3). Taking advantage of the open-well architecture of the device, we designed an experimental setup to decouple systemic endothelial priming from localized chemokine gradients. Systemic endothelial priming was achieved by adding TNF-α 24 hours prior to PBMC perfusion, while a localized chemoattractant gradient was established by adding CXCL12 (with or without TNF-α) directly into the gel chamber at the time of PBMC addition, enabling independent control of endothelial activation and PBMC recruitment across five experimental conditions (Fig. 3a).

**Figure 3.**
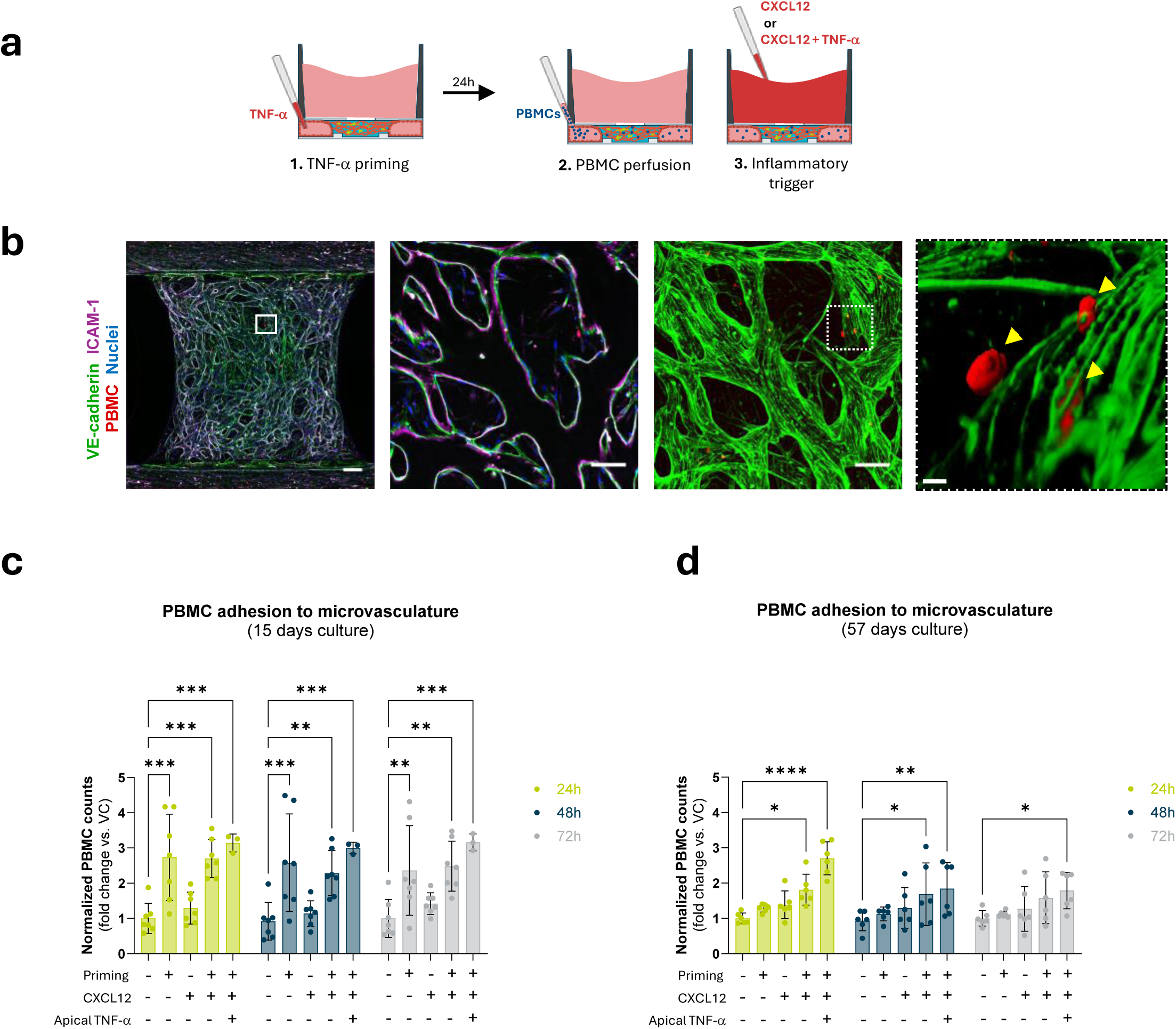
| Maturation-dependent immune cell adhesion in response to inflammation. a, Schematic of the experimental setup illustrating the conditions used to decouple systemic endothelial priming from localised chemokine gradients. Systemic priming was achieved by adding TNF-α to the perfusion channel 24 hours prior to PBMC perfusion. A localised chemoattractant gradient was established by adding CXCL12, with or without TNF-α, directly into the gel chamber at the moment of PBMC addition. Experiments were performed in both 15-day and 57-day cultures. **b,** Representative immunofluorescent images of a microvascular network subjected to TNF-α priming and PBMC perfusion with localised TNF-α and CXCL12 stimulation. After 72 hours, cultures were fixed and stained for ICAM-1 (magenta) and VE-cadherin (green). Left image shows the complete network acquired through a 4-site acquisition using a 10X objective; scale bar, 200 µm. The white square indicates the region shown at higher magnification in the adjacent image, representing a single z-plane acquired as a single-site with a 20X objective illustrating luminal ICAM-1 expression (magenta); scale bar, 100 µm. The third image represents the same region as a maximum intensity projection acquired with a 20X objective, showing VE-cadherin (green) and perfused PBMCs (red). The dotted white square indicates the region shown in the rightmost image, a 3D reconstruction illustrating the sequential stages of PBMC transmigration: luminal adhesion, transendothelial migration, and extravasation into the surrounding ECM; scale bar, 100 µm. **c,** VC-normalised fold change of PBMC adhesion at day 15 across four treatment conditions and VC at 24, 48, and 72 hours post-PBMC perfusion (N=2, n=7 chips per condition; N=1, n=3 for the condition with TNF-α in gel chamber). **d,** Same quantification at day 57 (N=2, n=6 chips per condition). Data are presented as mean ± SD. Two-way ANOVA revealed a significant effect of condition (c: P<0.0001; d: P<0.0001) with no significant effect of timepoint (c: P=0.647; d: P=0.050) or condition × timepoint interaction (c: P=0.992; d: P=0.431). Exact P values for all pairwise comparisons are provided in Supplementary Tables 8 and 9. Significance thresholds: *P<0.05, **P<0.01, ***P<0.001, \*\*\**P<0.0001*.

Prominent luminal expression of ICAM-1 along VE-cadherin-positive endothelial walls was observed in 15-day networks fixed 72 hours after systemic TNF-α priming and localised TNF-α and CXCL12 stimulation, confirming endothelial activation under the combined condition (Fig. 3b). To confirm that the platform supports the complete leukocyte extravasation cascade, we created high-resolution 3D reconstruction of TNF-α-primed networks following PBMC perfusion. A representative reconstruction provided visual evidence of all three stages of transmigration (yellow arrows): initial docking and adhesion to the luminal vessel wall, active transendothelial migration characterized by a PBMC intersecting the VE-cadherin junction, and extravasation into the surrounding ECM (Fig. 3b).

Quantification of PBMC adhesion at Day 15 revealed a hierarchical recruitment response across experimental conditions (Fig. 3c). Systemic TNF-α priming alone significantly increased adhesion relative to vehicle controls (P = 0.0002; 2way ANOVA), while a CXCL12 gradient in the absence of endothelial priming was insufficient to drive significant recruitment. The highest recruitment levels, approximately 3-fold above vehicle controls, required the synergistic combination of systemic TNF-α priming and localized CXCL12 and TNF-α in the graft chamber. Adhesion levels in singly primed networks declined progressively from 24 to 72 hours, while the combined stimuli condition sustained recruitment across the full observation period.

To determine whether the maturity of the microvasculature modulates immune responsiveness, we performed identical assays on long-term cultures (Fig. 3d). At Day 57, mature microvascular networks exhibited a distinct recruitment profile compared to Day 15 cultures. Systemic TNF-α priming alone produced no significant increase in PBMC adhesion at any measured timepoint. Significant recruitment was exclusively observed in the presence of a localized chemoattractant gradient, with the combined CXCL12 and TNF-α condition reaching fold-changes comparable to those observed at Day 15. Adhesion levels declined more rapidly in 57-day vessels, with only the combined stimuli condition maintaining a significant difference from control by 72 hours PBMC post-perfusion (P=0.0245). Collectively, these findings demonstrate that the platform supports PBMC recruitment, transmigration, and extravasation in response to inflammatory stimuli, and that the maturation state of the microvascular network modulates the magnitude and duration of the immune cell adhesion response.

### Multiparametric analysis of acute compound exposure reveals dose-dependent vascular-stromal toxicity modulated by vascular maturation

To evaluate the platform’s capacity to resolve compound-specific vascular toxicity profiles using scalable, image-based readouts, six compounds spanning three mechanistic categories were screened for acute vascular toxicity in day 15 microvascular cultures. These comprised multi-targeted tyrosine kinase inhibitors (TKIs; lenvatinib, sorafenib, and cabozantinib), intracellular process modulators (LY2090314, a GSK-3 inhibitor, and chloroquine, an autophagy inhibitor), and the non-selective cytotoxic agent doxorubicin. Responses were quantified across five complementary readouts: cytotoxicity (nuclear coverage), vascular perfusability, apparent permeability (Papp), stromal remodelling (αSMA), and inflammatory activation (ICAM-1), enabling multi-parametric characterization of compound-specific vascular toxicity profiles (Fig. 4a).

**Figure 4.**
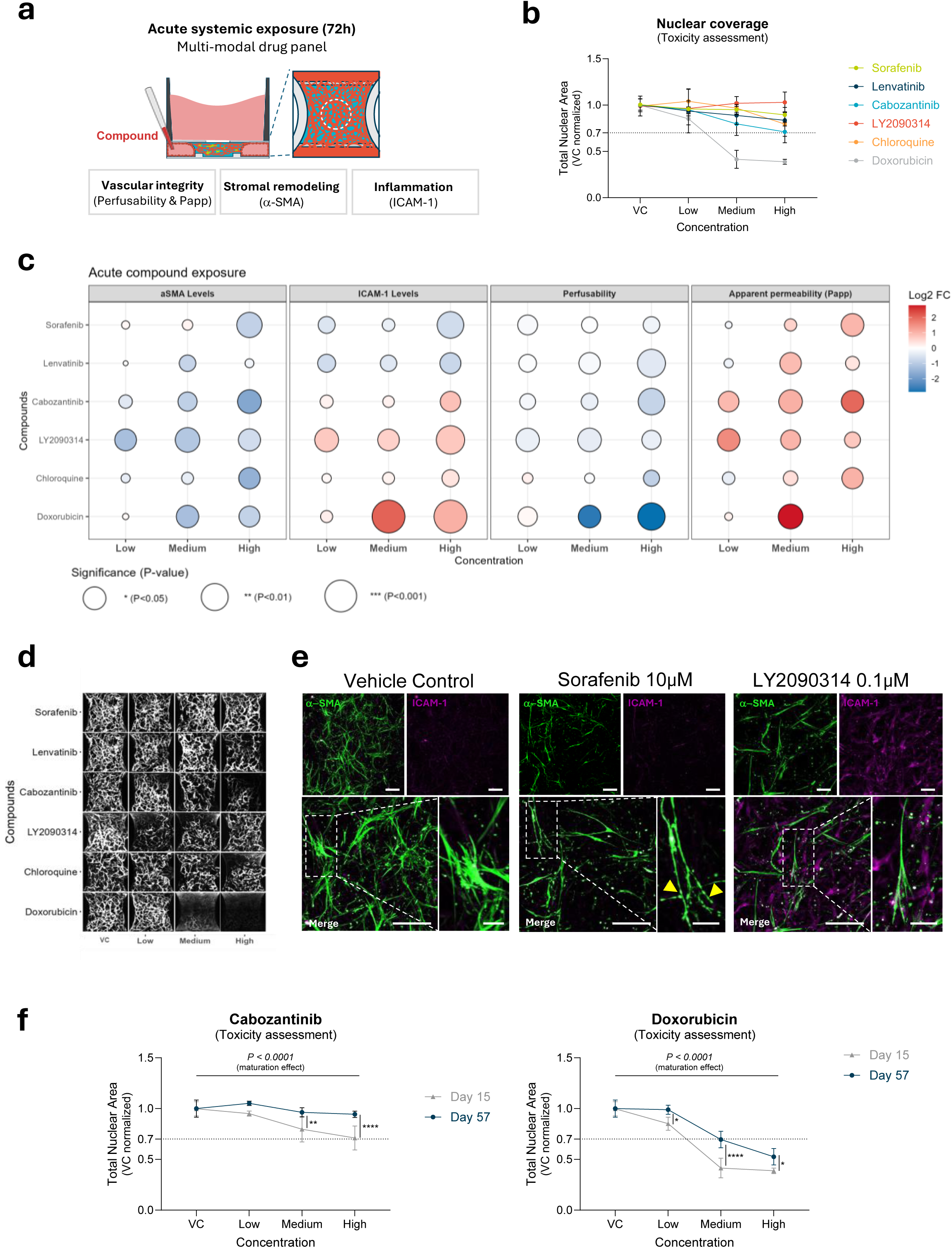
| Multiparametric analysis of acute compound exposure reveals dose-dependent vascular-stromal toxicity modulated by vascular maturation. a, Schematic of the experimental design illustrating acute (72-hour) exposure of microvascular networks to six compounds administered exclusively through the perfusion channel. Vascular toxicity was characterised across five readouts: nuclear coverage, αSMA and ICAM-1 expression levels, perfusable area fraction, and dextran extravasation fraction. **b,** Nuclear coverage as a measure of cytotoxicity following exposure to six compounds at four concentrations at day 15 (N=2 independent experiments across 12 plates, n=9 chips for VC n=6–7 chips per compound). Data are presented as mean ± SD. Two-way ANOVA revealed significant effects of compound (F(5,139)=41.30, P<0.0001), concentration (F(3,139)=47.49, P<0.0001), and a significant compound x concentration interaction (F(15,139)=10.70, P<0.0001). Dunnett’s post hoc test was applied relative to VC; exact P values for all pairwise comparisons are provided in Supplementary Table 10. Dashed line indicates the 70% viability threshold. **c,** Multiparametric bubble plot summarising compound-induced changes across all five readouts. Bubble colour represents log2 fold change relative to VC; bubble size represents statistical significance. All readouts were assessed per compound using Welch one-way ANOVA with Dunnett’s T3 post hoc test relative to VC (N=2 independent experiments, n=9 chips for VC, n=6–7 chips per compound). Exact P values for all comparisons are provided in Supplementary Table 11. **d,** Representative maximum intensity projection images of network perfusability for VC and each compound at low, medium, and high concentrations. Images represent 4-site acquisitions using a 10X objective. **e,** Representative immunofluorescent images of αSMA and ICAM-1 expression in VC, sorafenib (10 µM), and LY2090314 (0.1 µM) conditions. Images were acquired as single-site maximum intensity projections using a 10X objective; scale bar, 100 µm. Lower panels show digitally magnified insets from the corresponding upper images; scale bar, 100 µm (left) 25 µm (right). Yellow arrows indicate cytoplasmic vesicle accumulation in αSMA-positive cells following sorafenib exposure. **f,** Comparative quantification of total nuclear area following acute exposure to cabozantinib and doxorubicin in day 15 versus day 57 microvascular cultures (N=2, n=9 chips for VC, n=6–7 chips per compound). Data are presented as mean ± SD. Two-way ANOVA with Dunnett’s post hoc test relative to VC; exact P values are provided in Supplementary Table 12. Images in d and e are representative of the stated N and n. Significance thresholds: *P<0.05, **P<0.01, ***P<0.001, \*\*\**P<0.0001*.

Cytotoxicity screening revealed compound-specific sensitivity profiles across the panel (Fig. 4B). LY2090314 and Sorafenib showed no significant reduction in nuclear coverage across all concentrations (ns at all doses; two-way ANOVA with Dunnett’s post hoc test; n = 9), while lenvatinib, and chloroquine produced mild reductions only at their highest concentrations (P = 0.0029, and P = 0.0003, respectively), remaining above the 70% viability threshold in all cases. Cabozantinib produced a modest dose-dependent reduction approaching the viability threshold at medium and high concentrations (P < 0.0001), while doxorubicin was the only compound to reduce average nuclear coverage by more than 30% across all concentrations tested (low: P = 0.0123; medium and high: P < 0.0001), consistent with its non-selective mechanism of action.

Multi-parametric profiling revealed mechanistically distinct vascular toxicity signatures across compound classes (Fig. 4c). Among the TKIs, sorafenib produced a selective suppression of αSMA at high dose (P = 0.0077, Welch ANOVA with T3 Dunnett’s post hoc test; n = 6–7) without ICAM-1 increase, and mild to non-significant changes across Papp and perfusability, indicating a predominantly stromal effect. Lenvatinib produced a statistically significant increase in Papp at medium concentration (P=0.0393), with a marked perfusability decrease at the highest dose (P=0.0019), and no significant changes in αSMA, demonstrating that barrier disruption and perfusion loss can occur as independent vascular endpoints. Cabozantinib produced the broadest multi-parametric disruption among the TKIs, with progressive αSMA suppression, impaired perfusion, strong Papp increase, and concurrent ICAM-1 elevation.

LY2090314 exhibited a non-monotonic concentration-response relationship, with the lowest concentration producing the greatest αSMA reduction and largest Papp increase among the intracellular modulators, while ICAM-1 was elevated across all concentrations. The effects of chloroquine were generally mild and primarily observed at the highest concentration, where a simultaneous decline in vascular perfusion and α-SMA expression coincided with a statistically significant increase in Papp. Doxorubicin produced the most severe multi-parametric disruption across the entire panel, with perfusability collapse at medium and high concentrations, accompanied by marked ICAM-1 elevation (P < 0.0001), progressive αSMA loss, and Papp increase. Effects on network perfusion are also represented in Figure 4d.

Immunofluorescence imaging resolved two phenotypically distinct fibroblast responses across treatment conditions (Fig. 4e). In vehicle-treated cultures, αSMA-positive NHLFs displayed a highly branched morphology with extensive cytoplasmic projections. Both high-dose sorafenib and low-dose LY2090314 reduced αSMA area, yet produced distinct morphological outcomes: sorafenib-exposed fibroblasts displayed cytoplasmic vesicle accumulation (yellow arrows), whereas LY2090314-treated cells adopted a spindled, bipolar morphology with smooth, unbranched projections. Notably, LY2090314-induced ICAM-1 upregulation was localized predominantly to stromal fibroblasts within the ECM rather than to the vascular endothelium, a cell-type-specific distribution that would be unresolvable in conventional monolayer assays.

Comparison of Day 15 and Day 57 microvascular cultures exposed to cabozantinib and doxorubicin revealed compound-specific patterns of maturation-dependent pharmacological sensitivity (Fig. 4f). Cabozantinib-induced cytotoxicity was completely abrogated in Day 57 cultures, with nuclear coverage showing no significant reduction across any concentration while Day 15 cultures showed progressive reductions at medium and high doses (interaction P = 0.0006; two-way ANOVA with Tukey’s post hoc test; high dose Day 15 vs. Day 57: P < 0.0001). For doxorubicin, cytotoxicity was attenuated but not abolished in mature networks, with Day 57 cultures maintaining significantly higher viability than Day 15 counterparts at medium and high doses despite remaining below vehicle control levels (interaction P < 0.0001; medium and high dose Day 15 vs. Day 57: P < 0.0001 and P = 0.042, respectively).

Together, these results demonstrated that the platform resolves compound-specific, multi-parametric vascular toxicity profiles and that the maturation state of the microvascular network is a critical determinant of pharmacological sensitivity, ranging from complete abrogation to partial attenuation of cytotoxic responses.

### Chronic sorafenib exposure reveals progressive vascular remodelling and functional decline not detected during acute treatment

To determine whether the limited functional impact observed under acute sorafenib exposure reflected true vascular tolerance or simply an insufficient observation window, we conducted a 21-day chronic toxicity study leveraging the extended culture longevity of the platform. Microvascular networks were allowed to mature until Day 19 before chronic exposure to either 1 µM or 10 µM sorafenib until Day 40 (Fig. 5a).

**Figure 5.**
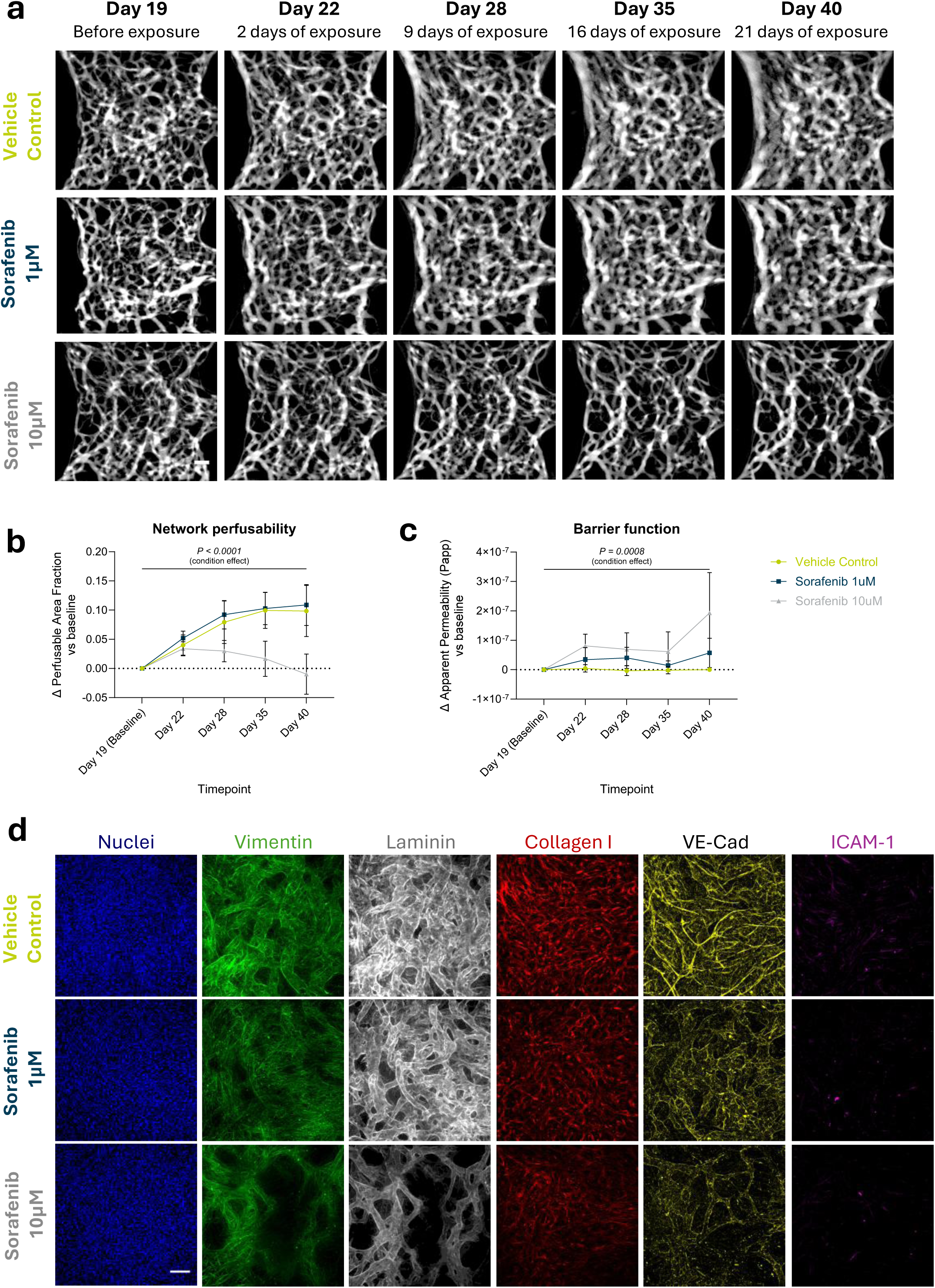
| Chronic sorafenib exposure progressively impacts vascular remodelling and function in a dose- and time-dependent manner. a, Representative fluorescence images of 70 kDa FITC-dextran perfusion assay performed at five timepoints across three exposure conditions. Single-site acquisition using a 4X objective; scale bar, 200 µm. ***b,*** Change in perfusable area fraction from baseline (Δ perfusable area fraction) across the chronic exposure window. Each data point represents an individual chip (N=1 experiment across 2 independent plates, n=6–8 chips per condition). Data are presented as mean ± SD. Mixed-effects model (REML) with Geisser-Greenhouse correction revealed significant effects of timepoint (F(1.891, 33.57)=49.74, P<0.0001), condition (F(2,18)=22.14, P<0.0001), and a significant timepoint × condition interaction (F(3.782, 33.57)=14.60, P<0.0001). Tukey’s post hoc test was applied for all pairwise comparisons; exact P values are provided in Supplementary Table 13. ***c,*** Change in Papp from baseline as a measure of vascular permeability across the chronic exposure window. Permeability was assessed using 70 kDa FITC-dextran (N=1 experiment across 2 independent plates, n=6–8 chips per condition). Data are presented as mean ± SD. Mixed-effects model (REML) with Geisser-Greenhouse correction revealed significant effects of time (F(1.307, 19.28)=10.13, P=0.0028), condition (F(2,16)=11.66, P=0.0008), and a significant time × condition interaction (F(2.614, 19.28)=4.949, P=0.0127). Tukey’s post hoc test was applied for all pairwise comparisons; exact P values are provided in Supplementary Table 14. **d,** Representative endpoint immunofluorescence images of microvascular networks following chronic sorafenib exposure. Staining for VE-cadherin (yellow), vimentin (green), laminin (grey), collagen I (red), and ICAM-1 (magenta). Maximum intensity projections of single-site images acquired with a 10X objective; scale bar, 100 µm. Images in a and d are representative of the stated N and n. Significance thresholds: *P<0.05, **P<0.01, ***P<0.001, \*\*\**P<0.0001*.

Perfusion assays revealed dose-dependent and temporally distinct functional trajectories that only emerged beyond the acute exposure window (Fig. 5b). While no significant differences were detected between conditions at Day 22, vehicle control and 1 µM sorafenib-treated networks both showed progressive increases in perfusable area fraction from baseline thereafter, whereas 10 µM sorafenib arrested this physiological expansion entirely, with perfusable area remaining near baseline through Day 35 before declining below baseline by Day 40, resulting in a significant and progressive divergence from Day 28 onward (VC vs. 10 µM at Day 40: P = 0.0019; 1 µM vs. 10 µM: P < 0.0001; mixed-effects model with Tukey’s post hoc test; n = 6–8 per condition). Vascular barrier function showed a mechanistically distinct and temporally offset response, with 1 µM sorafenib producing a consistent trend towards elevated permeability across multiple timepoints and 10 µM sorafenib driving significant barrier disruption at Day 22 and Day 40 (VC vs. 10 µM: P = 0.010 and P = 0.040, respectively), while vehicle control networks remained stable throughout (Fig. 5c).

Endpoint structural characterization revealed substantial remodelling at both concentrations (Fig. 5d). At 1 µM, a concentration that produced no detectable acute toxicity and maintained perfusable network density throughout the chronic period, VE-cadherin staining revealed a marked and selective elimination of narrow collateral branches, resulting in a simplified and streamlined vascular architecture not captured by perfusion metrics. Vimentin staining further revealed pronounced decoupling of NHLFs from the endothelial abluminal surface, disrupting the stabilizing perivascular architecture established during maturation. At 10 µM, these changes were severely exacerbated: fibroblast density was visibly reduced, Laminin expression markedly diminished, Collagen I deposition showed a dose-dependent decline, and VE-cadherin staining revealed loss of primary functional vessels consistent with the perfusion data. Across both doses, ICAM-1 expression remained undetected, demonstrating that this progressive structural toxicity occurs through a non-inflammatory mechanism consistent with acute exposure data.

Collectively, these findings demonstrate that the vascular MPS enable the detection of chronic, dose-dependent vascular toxicity that only emerges beyond acute exposure windows, capturing subclinical structural remodelling and perivascular disruption at therapeutically relevant concentrations.

## DISCUSSION

The transition toward New Approach Methodologies (NAMs) reflected in recent regulatory frameworks^27–29^ has increased the demand for predictive human vascular models capable of capturing clinically relevant drug responses over extended time periods. Although cardiovascular toxicity remains an important cause of late-stage drug attrition, most vascular MPS either simplify vascular architecture at the expense of physiological fidelity or achieve greater complexity with limited scalability, reproducibility, or longevity^11^. The microvascular model presented here addressed this gap through pump-free unidirectional perfusion of 3D microvascular networks for at least 57 days in a scalable 32-chip format, enabling both acute and chronic studies in mature vessels.

A persistent challenge in self-assembled vascular models is variability arising from stochastic network formation. A key element addressing this is the use of gravity driven flow, as this allows perfusion of 32 chips per plate and up to 16 plates per rocker platform to be perfused with exactly the same flow conditions, and without the risk of bubble trapping or need for extensive know how of microfluidic systems. Using variance decomposition^30^, we showed that system variability remains low across independent experiments, with intra-plate coefficients of variation below 10%. This suggests that variation is predominantly biological in origin rather than procedural, a distinction that is particularly important for predictive toxicology, where assay sensitivity depends on the ability to distinguish compound-induced phenotypes from intrinsic system noise. The low operator dependence observed further supports the workflow robustness that is essential for future regulatory qualification.

Together with architecture, culture time is a critical parameter required for the applicability of vascular MPS^11^. We address this by showing sustained 57-day vascular perfusion, demonstrating that cultures are not only viable, but are functional as well. Prior work has shown vascular plexi under bidirectional flow^31,32^ in which functional perfusion was time limited. Applying unidirectional flow through passive levelling of liquid through reciprocal rocking is a crucial innovation, keeping microvasculature accessible over prolonged periods of time and driving perivascular niche maturation. Through this approach, we achieved the longest reported culture span for perfusable SAMNs to date. Not only is this achieved exclusively through gravity-driven flow, but also using primary cells, a distinction from previous studies that employed actuator-driven^33^ or pump-based recirculation systems on cell lines ^25^.

Aspects including mechanical cues, microenvironmental complexity and flow characteristics have been implicated in vascular homeostasis and functionality^34–36^. Maturation of the vascular niche was witnessed by remodelling of lung fibroblasts vessel widening and collagen I deposition. NHLF progressively localized around endothelial structures in a manner suggestive of mural-like stabilization. Although multiple studies have described the role of lung fibroblasts in microvascular network formation and survival through paracrine signalling^37–39^, *in vitro* evidence of direct physical contact with endothelial vessels remains limited. Notably, Berthod et al. demonstrated that co-culture of HUVEC with dermal fibroblasts in a collagen sponge gave rise to αSMA^+^ pericyte-like cells around capillaries after 31 days^40^. This finding suggests that the brief culture periods typical of many *in vitro* culture systems may prevent the emergence of vascular-stromal dynamics that only arise in more mature microenvironments. Furthermore, concurrent accumulation of basement membrane proteins and stromal-derived collagen I suggests progressive ECM remodelling, potentially recapitulating the transition from provisional fibrin toward more stable, tissue-specific architectures. Despite passive flow, the network replicated the anisotropic vessel alignment and gradual luminal expansion reported in active-pump systems, with the latter suggested to be driven by growth factor gradients and evolving ECM stiffness^25,41^. Importantly, vessels remained within the physiological microvascular range (<100 µm), in contrast to the larger diameters observed under pump-driven recirculation^25^.

Structural maturation was accompanied by measurable functional changes. PBMC adhesion upon inflammatory stimulation was decreased in day-57 vessels, consistent with a more quiescence phenotype of the endothelium, an actively maintained state in mature vessels that raises the threshold for inflammatory activation^42,43^. This aligns with emerging evidence highlighting the role of ECM stiffness in leukocyte migration, where non-physiological stiffness promotes leukocyte attraction, chemotaxis, and adhesion ^44,45^. Consistently, endothelial cells on collagen I show reduced neutrophil transmigration compared to fibronectin^46^, suggesting that the progressive ECM remodelling observed here may modulate the inflammatory responsiveness at later culture stages. Additionally, glycocalyx maturation under sustained flow may further attenuate leukocyte adhesion by dampening pro-inflammatory signalling and limiting adhesion molecule expression^47,48^. Ultimately, the requirement for a localised CXCL12 gradient in mature networks suggests an increased dependence on microenvironmental cues as the vasculature matures. Whether reduced PBMC adhesion upon an inflammatory trigger directly derives from an elevated activation threshold, accelerated resolution of adhesion molecule expression, or a combination of both, is an aspect for future investigation.

The platform resolved acute compound-specific effects at the vascular and stromal levels. Within the vasculature, the lenvatinib-mediated decrease in perfusion aligns with pharmacovigilance data identifying its strong cardiovascular signal among VEGFR-TKIs, alongside clinical evidence of rapid endothelial dysfunction^49,50^. Interestingly, permeability remained unchanged, suggesting that vascular perfusion and barrier regulation are at least partially dissociable processes, consistent with TKI-mediated vascular normalization reported in the literature ^51^. Conversely, sorafenib’s greater reduction of αSMA-positive cells compared to lenvatinib reveals a distinct stromal vulnerability, suggesting this subpopulation relies heavily on RAF/MEK/ERK signalling over VEGFR/FGFR activation. Because TKI effects on healthy vascular stroma remain largely unevaluated, these insights provide critical context for off-target normal tissue toxicity. Importantly, this requirement for perivascular stability extends beyond kinase inhibition. Consistent with the pleiotropic role of GSK-3-mediated signalling in endothelial integrity^52,53^ and cytoskeletal regulation^54^, treatment with LY2090314 impaired perfusion and reduced α-SMA levels without overt cytotoxicity. This points to a disruption of perivascular contractile differentiation that aligns with the observed reduction in stromal branching.

The comparison between cabozantinib and doxorubicin responses further demonstrates that vascular maturation alters pharmacological vulnerability. Cabozantinib toxicity was markedly attenuated in mature networks, consistent with the established principle that mural cell-stabilised mature vessels are less susceptible to antiangiogenic agents than nascent vasculature^55,56^. This phenomenon has equally been implicated in tumour resistance to TKIs, where agents that demonstrated efficacy during early tumour growth show limited effect in tumours with established vasculature, an outcome attributable to changes in vascular phenotype and vessel stabilisation^57,58^. Doxorubicin responses were similarly attenuated but not abolished, suggesting partial protection associated with reduced endothelial proliferation rather than complete resistance. Together, these findings highlight the importance of matured vascular networks for studying vascular toxicity.

The capacity of the platform to reproduce concentration- and time-dependent vascular phenotypes under chronic sorafenib exposure reflects a clinically relevant toxicity spectrum. Hypertension is a common off-target effect of VEGFR-targeting TKIs, with sorafenib associated with all-grade incidences of approximately 19–23% across randomised clinical trials^59^. Mechanistically, VEGFR inhibition has been linked to impaired eNOS/NO signalling and progressive microvascular rarefaction, proposed drivers of the observed blood pressure elevation^60^. In the present system, prolonged sorafenib exposure reproduced several features of this phenotype spectrum. The hierarchical susceptibility of smaller vessels aligns with the VEGF-dependence of capillary-level maintenance signalling^61^, while the temporal precedence of perivascular NHLF decoupling over perfusion loss is functionally reminiscent of the pericyte destabilisation described following PDGFR-axis disruption by other TKIs^62^. At higher concentrations, the abrogation of vessel area increase and ultimate loss of perfusable area reflect the anti-proliferative and pro-regressive effects of sorafenib, ultimately leading to vascular rarefaction. The progressive, concentration-dependent emergence of severe barrier dysfunction parallels clinical observations in which renal vascular injury may occur independently of systemic hypertension^63^. Importantly, these structural alterations occurred without detectable ICAM-1 induction in mature microvascular networks, supporting a model in which VEGFR-TKI vascular toxicity is driven primarily by impaired survival signalling and progressive structural remodelling rather than classical inflammatory endothelial activation.

More broadly, the divergence between acute and chronic responses underscores an important gap in current vascular toxicology practice. Clinically relevant TKI-associated phenotypes, including rarefaction, metabolic adaptation, and persistent vascular remodelling, emerge progressively during prolonged exposure and are therefore poorly represented in short-term assays. Increasing evidence further suggests that chronic kinase inhibition induces metabolic and epigenetic reprogramming before stable pathological states are established, often also associated with development of treatment resistance^64^. The extended culture window we presented here is therefore valuable not only because it complements existing assays, but because it enables experimental access to delayed vascular biology and chronic adaptive states that are increasingly relevant to emerging regulatory toxicology frameworks.

Several limitations of the present study should be acknowledged. Although flow within the platform is predominantly unidirectional, gravity-driven perfusion through rocking inherently includes brief intervals of backflow, up the 8 seconds in every 80 second cycle. However, the limited functional impact of this phenomenon is supported by sustained endothelial alignment and prolonged vascular stability. The use of HUVECs does not capture organ-specific endothelial heterogeneity; however, standardised commercial cell sources were intentionally selected to minimise donor-dependent variability and facilitate reproducibility assessment. Incorporation of tissue-specific endothelial populations and dedicated pericytes will likely improve physiological fidelity in future work. Maturation-dependent functional responses were inferred from phenotypic observations rather than direct molecular analysis; future studies should more comprehensively characterise glycocalyx composition, adhesion molecule kinetics, and matrix properties at successive timepoints. Following the current demonstration of the model’s potential, a broader chronic pharmacological benchmarking against clinical datasets, will be an important next step toward regulatory qualification of the platform.

Grounded in image-based approaches, this study builds on a series of phenotypic readouts that are non-invasive and highly scalable. Collectively, these form a practical, comprehensive toolkit capable of extracting meaningful time resolved toxicological information, supporting standardised implementation across both academic and industrial settings. Beyond vascular toxicology, the modular architecture of the stable microvascular model in MPS provides a foundation for vascularised organ models allowing for the investigation of epithelia-stroma interactions through integration of epithelial and tissue-specific compartments, with immediate relevance to lung, gut, and related barrier tissues.

In conclusion, this work demonstrates that long-term vascular homeostasis and progressive maturation are defining aspects in the future of vascular MPS and that these can emerge *in vitro* without external pumps, lowering the barrier to entry, simplifying operation and integrating with standard laboratory equipment. With growing regulatory expectations for NAMs, formalized through qualification programs such as FDA’s Innovative Science and Technology Approaches for New Drugs (ISTAND)^28^ and EMA’s Qualification of Novel Methodologies (QoNM)^29^, this advancement enhances the position of vascular MPS as a predictive tool for human physiological and toxicological responses.

## METHODS

### 2D Cell culture

Primary Human Umbilical Vein Endothelial Cells (HUVECs; LONZA, C2519A,) were expanded in T175 culture flasks (Nunc™ EasyFlask™, FischerScientific, 10246131) using EGM-2 endothelial medium (LONZA, CC-3162). Primary Normal Human Lung Fibroblasts (NHLFs; LONZA, CC-2512) were expanded in T75 flasks (Corning, 430641U) and cultured using FGM-2 medium (LONZA, CC-3132). Both cell types were used between passages 4–5. All cultures were maintained at 37 °C, 5% CO₂ under humidified conditions and regularly screened for mycoplasma (LONZA, LT07-710). Cells were harvested after 4 days in culture and dissociated according to the manufacturers’ protocols prior to seeding into the OrganoPlate.

### OrganoPlate Graft 32 UF

The OrganoPlate Graft 32 UF (MIMETAS, 3203-400-E) consists of 32 microfluidic chips embedded within a 384-well plate footprint. Each microfluidic chip comprises a central gel chamber with a 1 mm diameter access hole where cells within an extracellular matrix (ECM) gel can be seeded, a perfusion channel, and a bypass channel. Phaseguides^TM^ – capillary pressure barriers patterned between the el chamber and perfusion channel^65^– allow precise ECM localization without physical membranes. Each chip comprises 2 long wells and 6 individual wells, the 384-well plate footprint was modified to form a pair of long wells for each microfluidic chip formed by connecting 3 wells together resulting in a 256-well plate (Fig.1a, b, wells 1-3 and 4-6). When placed at a high angle (25 °), the long wells, combined with a low media volume created an air liquid interface above the outermost inlet holes. This disconnects the fluid, preventing backflow by the surface tension of the fluid above the exposed interface, thereby creating a Laplace valve which prevents the channel from emptying. In the positive incline, flow is routed through the perfusion channel creating a pressure difference across the gel chamber, which in turn drives flow through the vascular network once it has formed connection across the gel chamber. During the negative incline fluid is redirected through the bypass channel and the system is reset. The design provides an active phase of 60 seconds with a reset phase of 20 seconds allowing for unidirectional flow which enables long-term, pump-free perfusion under physiologically relevant conditions (Fig. 1c,d).

### 3D Microvascular Culture in the OrganoPlate

HUVECs and NHLFs were embedded in a fibrin-based ECM. The ECM solution was prepared with 5 mg/mL human fibrinogen (Enzyme Research Laboratories, FIB 1) and 0.2 U/mL thrombin (Enzyme Research Laboratories, HT1002a). HUVECs and NHLFs were added at final concentrations of 10,000 cells/μL and 1,667 cells/μL, respectively. A 1.2 μL aliquot of the cell–ECM suspension was loaded into each gel chamber (Fig 1b, well B5) and polymerized for 15 minutes at 37 °C. Following polymerization, 50 μL of MV-2 medium (PromoCell, C-22022) was added to each gel chamber.

Perfusion channels were precoated with 30 μg/mL fibronectin (Sigma-Aldrich, F1141) for 1 hour at 37 °C, and seeded with HUVECs (10,000 cells/μL) by passive pumping as previously described^66^. After a 2-hour adhesion period, 100 μL of EGM-2 medium was added to the long wells and 50 μL to the gel chambers. Plates were placed on an OrganoFlow® rocker (MIMETAS, MI-OFPR-L) to establish perfusion. During the first 24 hours, bi-directional flow was applied (7° inclination, 8-min rocking interval). Perfusion was then switched to unidirectional flow (25° inclination, 1-min rocking interval to the left (Fig. 1C) and 20 seconds to the right side (Fig. 1D)). From day 1 to day 7, EGM-2 medium in the perfusion lanes was supplemented with 50 ng/mL VEGF (PeproTech, 100-20), 250 nM S1P (Sigma-Aldrich, 73914), and 100 KIU/mL aprotinin (Nordic Pharma, 7005124), while gel chambers received EGM-2 supplemented with aprotinin only. VEGF and S1P supplementation were discontinued after day 7. Medium was exchanged three times per week.

### Dye perfusion assay

To confirm vascular perfusability, 40 μL of EGM-2 medium containing 0.25 mg/mL FITC-dextran (150 kDa or 70kDa; Sigma-Aldrich, 46946 or 46945) was added to perfusion inlets and outlets, leaving gel chambers empty. Plates were placed on the OrganoFlow rocker (25° inclination, with a 1-min interval to the left followed by a 20-s interval to the right) for 4 minutes to facilitate perfusion. Fluorescence images were captured at 37 °C using the ImageXpress Micro XLS-C (Molecular Devices) with a 4X objective.

### Immune adhesion assay

To evaluate immune cell adhesion, 14 and 56-day microvascular networks were primed for 24 hours by adding medium supplemented with 2.25 ng/mL TNF-α (ImmunoTools, 11343015) exclusively to the perfusion channel. 24 hours post-priming, inflammatory triggers, consisting of 2.25ng/mL TNF-a and/or 800 ng/mL CXCL12 (PeproTech, 300-28A) diluted in EGM-2, were added to the gel chamber to establish a chemotactic gradient. Immediately after, single donor human Peripheral blood mononuclear cells (PBMCs; LONZA, CC-2702P) were stained with Incucyte® Cytolight Rapid Dye (Sartorius, 4705) according to the manufacturer’s protocol and resuspended in EGM-2 medium before they were introduced into the perfusion lane (50,000 cells/chip) and allowed to circulate through the microvasculature.

Following PBMC addition, plates were returned to the OrganoFlow and maintained under standard flow conditions. PBMC adhesion to the endothelium was assessed after 24, 48 and 72 hours using an ImageXpress Micro XLS-C high-content confocal system (Molecular Devices) with a 4X objective. The total number of adherent PBMCs within the vascular networks was quantified using Fiji and compared across experimental conditions. Following the 72-hour time point, cultures were fixed and processed for immunofluorescence as described in the "Fixation and Immunostaining" section.

### Compound exposure

Lenvatinib (TargetMol, T0520), sorafenib (MedChem Express, HY-10201), cabozantinib (Selleck Chemicals, S1119), LY2090314 (Selleck Chemicals, S7063), Chloroquine diphosphate (Selleck Chemicals, S4157), and Doxorubicin hydrochloride (MedChem Express, HY-15142) were prepared according to the manufacturer’s instructions and diluted in EGM-2 medium immediately prior to use.

Compounds were administered at day 15 of culture, exclusively via the perfusion channel (luminal perfusion), at three concentrations: Sorafenib, Lenvatinib, Cabozantinib, and LY2090314 (0.1, 1, 10 µM); Chloroquine (1, 50, 100 µM); and Doxorubicin (1, 5, 10 µM). Cabozantinib and Doxorubicin were additionally tested in cultures at day 57 using the same exposure conditions. Vehicle controls were included at matched solvent concentrations. Cultures were exposed for 72 hours, after which functional endpoints (microvascular perfusability and apparent permeability, Papp) were performed and the cultures were fixed for immunostaining.

For chronic exposure, microvascular networks were maintained until day 19 prior to treatment initiation. At that timepoint, sorafenib (1 and 10 µM) was administered exclusively via luminal perfusion for 21 days with compound-supplemented medium refreshed three times per week (every 48–72 h) to ensure consistent pharmacological exposure. Vehicle-treated controls were maintained in parallel under identical perfusion conditions. Vascular functionality, including perfusability and permeability, was monitored throughout the exposure window. At the experimental endpoint on day 40, cultures were fixed and processed for immunofluorescence staining.

### Fixation and Immunostaining

Cultures were fixed with 3.7% formaldehyde (Sigma-Aldrich, 252549) for 15 minutes. After fixation, samples were permeabilized and blocked for 2 hours in PBS containing 1% Triton X-100 (Sigma, T8787) and 3% bovine serum albumin (BSA; Sigma, A2153). Primary antibodies were diluted in incubation buffer (0.3% Triton X-100, 3% BSA in PBS) and applied overnight at room temperature. The following primary antibodies were used: mouse anti-PECAM-1 (DAKO, M0823), rabbit anti-Vimentin (Novus Biologicals, NBP1-31327), Human alpha-Smooth Muscle Actin (αSMA) Alexa Fluor® 488-conjugated Antibody (R&D Systems, IC1420G), mouse anti-ICAM-1 (R&D Systems, BBA3), rabbit anti-Fibronectin (Abcam, ab45688), rabbit anti-Laminin (Abcam, ab11575), Recombinant Alexa Fluor® 647 Anti-Collagen I antibody (Abcam, ab280968), rabbit anti-VE Cadherin (Abcam, ab33168).

After three washes in PBS with 0.3% Triton X-100, secondary antibodies and Hoechst 33342 nuclear stain (Thermo Fisher Scientific, H3570) were applied overnight at room temperature. Secondary antibodies included donkey anti-mouse Alexa Fluor 488 (Thermo Fisher, A21202), donkey anti-rabbit Alexa Fluor 647 (Abcam, ab150075), donkey anti-mouse Alexa Fluor 555 (Thermo Fisher, A31570) and goat anti-rabbit Alexa Fluor 555 (Thermo Fisher, A32732). Chips were washed 3 times with PBS + 0.3% Triton X-100 and imaged using the ImageXpress® Confocal HT.ai High-Content Imaging System (Molecular Devices).

Phenotypic changes were evaluated by immunofluorescence imaging of the full gel chamber (10x objective, 4 sites). αSMA expression was quantified as signal density within the stromal compartment using LabKit plugin (Fiji). ICAM-1 expression was quantified as integrated fluorescence intensity across the entire culture following global Otsu thresholding.

Total nuclear area was utilized as a sensitive, high-throughput proxy for cytotoxicity, as it allows for the objective, automated quantification of morphological hallmarks of cell death such as chromatin condensation and nuclear fragmentation without the reliance on membrane-integrity-based enzymatic assays^67,68^. All image-based analyses were performed in Fiji (ImageJ) using consistent segmentation models and thresholding parameters across all experimental conditions.

### Analysis of vascular network morphology and integrity

Images of the dye perfusion assay were pre-processed using Fiji to reduce background signal by applying a rolling ball background correction routine^69,70^. The centre of the gel chamber was selected and segmented using a trainable classifier in LABKIT as previously reported^24,71^ .Vessel signal was extracted as foreground (white), and non-vessel signal was considered background (black). The segmented vessel signal was then skeletonized, after which vascular characteristics were extracted. Extracted values were then compared across culture timepoints.

Vessel orientation was quantified from pre-processed perfusion images using the Directionality plugin in Fiji (v1.54f). For each microvascular network at two distinct timepoints (day 8 and 57), the dominant orientation (Direction) was determined by a Gaussian fit of the orientation histogram. The alignment prevalence (Amount) was calculated as the normalized sum of the raw histogram values within one standard deviation of the Gaussian centre. These parameters were visualized in R using rose plots, where the angular position and radial length of each needle represent the Direction and Amount, respectively, for individual networks.

Vascular network integrity was assessed by calculating the dextran retention fraction (DRF) in networks from three different plates across six culture timepoints (Supplementary methods).

### Network perfusability and permeability of compound exposure

Microvascular perfusability was assessed at baseline (immediately prior to treatment) and 72 h post-compound exposure as previously described in “Dye Perfusion Assay” section. A 70 kDa FITC-Dextran (Sigma-Aldrich, 46945) was introduced into the perfusion channels to visualize the patent vascular network. Imaging was conducted using an ImageXpress HT.ai high-content imaging system (Molecular Devices) equipped with a 10× objective. For each chip, four overlapping sites spanning the entire gel chamber were acquired and subsequently stitched to generate a comprehensive representation of the microvascular bed. Automated network segmentation was performed using IN Carta software (Molecular Devices, version 2.8). Changes in functional perfusability were quantified as the difference in the total perfusable area between the initial and final time points. Vessel Papp was quantified according to previously described methods ^72,73^. A more detailed explanation of the methodology can be found in the supplementary methods.

### Statistical Analysis

Data analysis was performed in GraphPad Prism (version 10). Normality was assessed using the Shapiro–Wilk test prior to all analyses. Operator effect was assessed using nested one-way ANOVA, with individual chip values nested within experiments to account for the hierarchical data structure. Longitudinal morphological readouts including vessel integrity, perfusable area fraction, and vessel width were analyzed using repeated measures one-way ANOVA with Dunnett’s post hoc test. PBMC adhesion was evaluated using two-way ANOVA with Dunnett’s post hoc test to assess the effects of timepoint and condition. For acute compound exposure, data were log2-transformed prior to analysis to ensure normality, confirmed by Shapiro–Wilk test. Welch ANOVA with Dunnett’s T3 post hoc test was applied across all compounds and readouts to ensure comparability.

Non-normally distributed datasets were analysed using Kruskal–Wallis with Dunn’s multiple comparisons test. Significance thresholds were set as follows: P<0.05 (*), P<0.01 (**), P<0.001 (***), P<0.0001 (****). Where the number of replicates is stated, N=1 refers to a single experiment comprised of one or more plates, n=1 refers to a single chip within a plate.

## ACKNOWLEDGEMENTS

The authors thank Aleksandra Olczyk for support with data analysis, Anastasia Kondratyeva and Tomás Coimbra for experimental support, Alexis Dalaud for chip fabrication, Frederik Schavemaker for assistance with schematic design, and Dr Henriëtte Lanz for valuable input

## DATA AVAILABILITY

The datasets used and/or analysed during the current study are available from the corresponding author on reasonable request.

## DECLRATIONS

### Ethics approval and consent to participate

Not applicable.

### Consent for publication

Not applicable.

