## Supplement. materials for "Scalable microvascular networks-on-chip enable 8-week unidirectional perfusion for long-term vascular toxicity screening"

### Supplementary figures

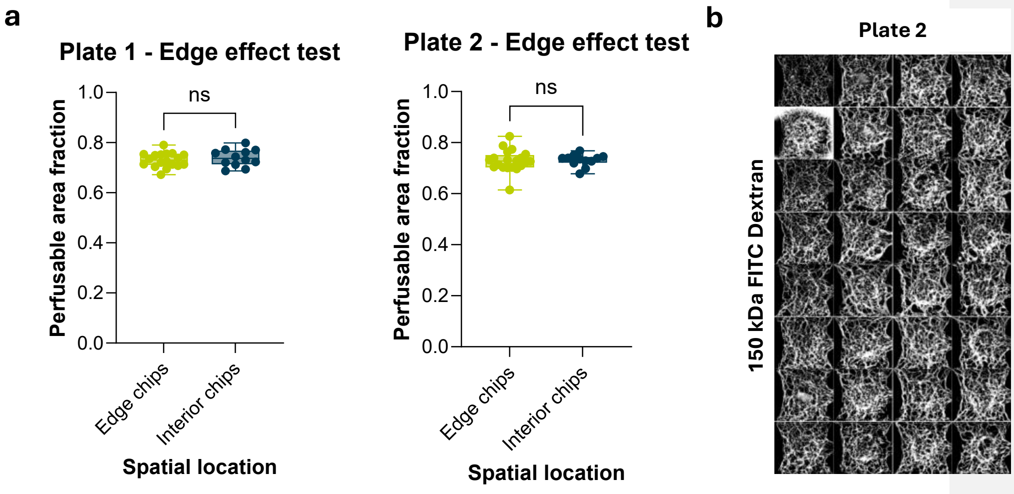


**Supplementary Fig. 1 | Intra-plate variability and edge effect. a,** Perfusable area fraction compared between edge and interior chips in two independent plates. Each data point represents an individual chip (Plate 1: n=18 edge, n=12 interior chips; Plate 2: n=20 edge, n=12 interior chips). Data are presented as mean ± SD. No significant difference in perfusable area fraction was detected between edge and interior chips in either plate (Plate 1: t(20.74)=0.730, P=0.474; Plate 2: t(29.94)=0.174, P=0.863; two-tailed unpaired t-test with Welch's correction). **b,** Representative fluorescence images of 150 kDa FITC-dextran perfusion assay across all chips of Plate 2 (20-day culture), demonstrating homogeneous microvascular network formation throughout the plate. Images acquired with a 4X objective.

### Supplementary methods

#### Dextran retention fraction (DRF)

By perfusing networks with 150 kDa FITC-dextran and imaging after a 4-minute incubation period. The high molecular weight of the tracer ensures that fluorescence leakage reflects paracellular barrier permeability rather than passive diffusion. Fluorescence images were acquired as maximum intensity projections under identical imaging conditions across all timepoints. A DRF was calculated as the fraction of total fluorescence signal detected in the extravascular compartment relative to the combined intra- and extravascular signal:

$$DRF = \frac{I^{total\ outside}}{I^{total\ vasculature} + I^{total\ outside}}$$

This ratio represents the degree to which dye was able to enter the surrounding ECM, filling up the local space until equilibrium is reached. Very leaky vasculature will have a high ratio, indicating that the inside-vasculature signal is close to the outside-vasculature signal, reaching 1 if inside and outside intensities are equal. Conversely, a leak tight vasculature will have values closer to 0.

#### Apparent permeability (Papp)

Apparent permeability was determined by monitoring the interstitial translocation of a 70 kDa FITC-Dextran tracer from the vascular lumen into the adjacent ECM compartment. A working solution of 0.25 mg/mL FITC-dextran was added to the long wells (150 µL inlet; 110 µL outlet). Following a 1-min incubation on the OrganoFlow rocker, time-lapse imaging was performed on the ImageXpress HTai using a 10× objective and Z-stacks with 8-µm steps. The Papp was calculated from maximum intensity projections based on the principle of mass conservation, according to the following equation:

$$Papp = \left(\frac{1}{I_V^{t1} - I_T^{t1}}\right) * \left(\frac{(I_T^{t2} - I_T^{t1})}{\Delta t}\right) * \left(\frac{A_{lateral_{tissue}}}{P}\right)$$

$I_V^{t1}$ and $I_T^{t1}$ represent the mean fluorescence intensities in the vascular and tissue regions, respectively, at the initial time point (t1). $I_T^{t2}$ is the mean tissue intensity at the second time point (t2). Δt is the measured time interval between image acquisitions. $A_{lateral_{tissue}}$ and $P$ are the 2D projected lateral tissue area and the vascular perimeter, respectively. The ratio $\frac{A_{lateral_{tissue}}}{P}$ was utilized to approximate the 3D volume-to-surface-area ratio $\frac{V_{tissue}}{A_{surface}}$ of the microvascular network within the 2D projected space. All intensity and geometric parameters were quantified using automated segmentation protocols in Fiji (version 1.54).

**Commented [TO1]:** You could opt to reference the AIM Biotech method paper here, instead of explaining the method.

**Commented [AR2R1]:** @Lenie van den Broek | MIMETAS @Todd Burton | MIMETAS I saw that the original protocol comes from these publications: Curry et al., 1983; Campisi et al., 2018. Should I just reference them and move it to supplements, or remove it entirely?

**Commented [TM3R1]:** move to suppliments and reference
